# The Bacteriocin Sublancin Restores Vancomycin efficacy against vancomycin-resistant enterococci *in vitro* and *in vivo*

**DOI:** 10.64898/2026.02.23.707390

**Authors:** Xinyu Guo, Yuzhe Zhao, Chenglong Li, Yage Sun, Yuan fei Lu, Yanling Wang, Yangke Liu, Bing Ma, Xiang-Dang Du, Dexi Li

## Abstract

Vancomycin-resistant enterococci (VRE) is one of the serious threat to global public health, with the diminishing effectiveness of antibiotics. There is an urgent need for novel strategies to control this multidrug-resistant bacterial infections. Here, our findings demonstrate that sublancin, a bacteriocin produced by *Bacillus subtilis,* exhibits intrinsic antibacterial activity and, more importantly, potentiates vancomycin, therapy restoring its efficacy against VRE. Using a series of *in vitro* assays, including fractional inhibitory concentration index, time-killing analysis, and resistance development assays, we show that sublancin significantly enhances the bactericidal effectiveness of vancomycin. *In vivo*, the sublancin-vancomycin combination therapy markedly reduced bacterial loads, improved svrvival rates in a *Galleria mellonella* model and enhanced bacterial clearance rates in a mouse model. Mechanistic studies using RT-PCR revealed that sublancin down-regulates the expression of the *vanA* resistance gene cluster. All in all, these findings make vancomycin and this antibiotic peptide combination as a promising candidatese to enhance vancomycin efficacy and overcome VRE infections, potentially through inteference with resistance gene expression.

## Introduction

Vancomycin-resistant enterococci (VRE) and staphylococci represent a major therapeutic challenge among gram-positive bacteria in clinical settings (1–4). The prevalence and transmission of these organism, mainly VRE, across human(1–4), animal husbandry (3, 5), and the environment reservoirs (6), cause a serious public health threat, contributing to increasing global incidence and substantial economic burden. The diminished efficacy of vancomycin-a traditional last-line therapy primarily results from the dissemination of resistance genes, such as *vanA* and *vanB* among gram-positive bacteria(1–9). Addressing this challenge requires a comprehensive One Health approach for effective control.

Daptomycin and linezolid, remain critical antibacterial agents for treatment for VRE (10, 11). However, the effect is increasingly limited by emerging resistance, or co-resistance. This therapeutic impasse underscores a critical gap in our antimicrobial arsenal and highlights the need for novel agents that can directly counter established resistance mechanism. Promising strategy to combat these infections included the development of next-generation vancomycin analogues (12), and the exploration of antimicrobial peptides (AMPs) or bacteriocins that either modulate host immunity to combat infections (13–15), or exert direct antibacterial activity (16–20), or target regulatory pathway such as transcription factor (21, 22).

In our preliminary routine screenings, a naturally bacterially derived AMP, sublancin (23), demonstrated an unexpected capacity to resensitize VRE to vancomycin. To further investigate this phenomenon, we evaluated the *in vitro* and *in vivo* activity of sublancin in combination with vancomycin by determining the minimum inhibitory concentrations (MICs) against clinical VRE isolates and elucidating the underlying mechanism. Collectively, this work provides a foundational basis for using bacterial AMPs to effectively counteract vancomycin resistance, offering a promising combinatorial strategy against VRE infections.

## MATERIALS AND METHODS

### Bacterial strains and regeants

#### Strains

*vanA*-positive VRE HN1, MRSA 1, *poxtA and fexB* positive *E. faecalis* E035, E076 were utilized for assessing antibiotic resistance reversal activity and investigating molecular mechanisms. All strains were obtained from the College of Veterinary Medicine at Henan Agricultural University. Species identification was confirmed by whole-genome sequencing and 16S rRNA analysis. *Enterococcus faecalis* JH2-2 and *Staphylococcus aureus* ATCC 29213 set as the control strains.

#### Regeants

Vancomycin, cefoxitin, florfenicol, tetracycline and erythromycin was purchased from Shanghai Macklin Biochemical Co., Ltd., and the antimicrobial peptide sublancin was obtained from Sinagri YingTai Bio-peptide Co., Ltd. The following reagents and kits were used: BCECF-AM fluorescent probe (AAT Bioquest), propidium iodide (Solarbio), *N*-phenylnaphthylamine (NPN, Yuanye Bio-Technology), ATP detection kit (Beyotime), ethidium bromide (EtBr) and carbonyl cyanide m-chlorophenylhydrazone (CCCP) from MedChemExpress, DiSC_3_(5) fluorescent probe, and the TRIzol Universal RT-PCR kit (Jiangsu ComWin Biotech).

### Antibacterial susceptibility test and microdilution checkerboard methods

Minimal inhibitory concentrations (MICs) of antibiotic and sublancin were individually determined using the broth microdilution method, following Clinical and Laboratory Standards Institute (CLSI) guidelines. Poential synergistic interactions betweeen selected antibiotic (vancomycin, cefoxitin, florfenicol, tetracycline and erythromycin) and sublancin were assessed via checkerboard assay. *E. faecalis* JH2-2, and *S. aureus* ATCC 29213 set as control strains.

For the fractional inhibitory concentration (FIC) assay, vancomycin and other test antibiotic with varying concentrations of sublancin against the *vanA*-positive VRE strain HN1, MRSA 1 and *E. faecalis* strains E035 and E076. The FIC index (FICI) was calculated as follows: ΣFIC = (MIC of drug A in combination / MIC of drug A alone) + (MIC of drug B in combination / MIC of drug B alone). The interaction was interpreted as synergistic if the FICI was ≤0.5, indifferent if >0.5 to ≤4.0, and antagonistic if >4.0.

### Time-Kill Assay

Time-kill assays were performed to evaluate the bactericidal activity of vancomycin alone and in combination with sublancin against VRE HN1. Mid-log phase bacterial cultures (∼1 ×10⁶ CFU/mL) were treated with vancomycin (¼×MIC), sublancin (¼×, or×^1^/_8_ MIC), sublancin (¼×, or×^1^/_8_ MIC) in combination with vancomycin (80µg/mL), or PBS as a vehicle control. Samples were collected at 0, 2, 4, 6, 8, 12, and 24 h, serially diluted at rato 1:10, and plated on Brain Heart Infusion agar (BHA). Viable colonies were enumerated after overnight incubation at 37°C.

### Development of Resistance Assay

The potential for resistance development was assessed using VRE strain HN1. The strain was diluted 1:1000 into fresh Brain Heart Infusion (BHI) broth containing subinhibitory concentrations (¼× MIC) of the test agents: vancomycin alone or vancomycin in combinnation with sublancin. Cultures were passaged daily for 32 days following 18 h of incubation at 37°C. The MIC was determined every fourth passage to monitor the changes in the susceptibility of the strain to vancomycin.

### Assessment of key indices of bacterial physiological state and energy metabolism

VRE strain HN1 was harvested during the logarithmic growth phase (OD_600_ = 0.5–0.8). Bacteria from five treatment groups were analyzed: an untreated control and groups treated with four concentrations of sublancin (20, 40, 80, and 160 µg/mL). Bacterial cells were pelleted by centrifugation at 3000 × g for 5 min, washed twice with phosphate-buffered saline (PBS), and resuspended to an OD600 of 0.5.

The following fluorescent probes were used at the indicated final concentrations and incubation conditions: 1-*N*-phenylnaphthylamine (NPN; 15 µM, 30 min, 37°C) to assess outer membrane permeability, propidium iodide (PI; 0.5 µM, 30 min, 37°C) to evaluate cytoplasmic membrane integrity, 3,3 ′ -dipropylthiadicarbocyanine iodide [DiSC3(5), 10 µM, 30 min, 37°C] to monitor membrane depolarization, and 2′,7′-bis-(2-carboxyethyl)-5-(and-6)-carboxyfluorescein acetoxymethyl ester (BCECF-AM, 10 µM, 30 min, 37°C) to measure the proton motive force (PMF). Cellular ATP levels and total reactive oxygen species (ROS) were quantified using commercial assay kits according to the manufacturer’s protocols, as previously described (24, 25).

For all fluorescence-based assays, 200-µL aliquots from each sample (n = 3 independent replicates per group) were transferred to a black 96-well plate, and fluorescence intensity was measured using a multifunctional microplate reader.

### Analysis of th**e** *vanA* operon expression by RT-qPCR

Total Bacterial RNA was extracted from VRE strain HN1 following four treatments: (i) an untreated control, (ii) sublancin alone, (iii) vancomycin alone, and (iv) combination of vancomycin and sublancin. The transcript levels of the vancomycin resistance genes *vanR*, *vanS*, *vanH*, *vanA*, *vanX*, *vanY* and *vanZ*, were quantified by reverse transcription quantitative PCR (RT-qPCR) (24, 26). Partial primer pairs were designed using Primer-BLAST (NCBI) with the following criteria: 18–22 bp in length and a melting temperature (Tm) of 60 ± 2°C. Amplification was performed using SYBR Green assays on Bio-Rad CFX96, with the 16S rRNA serving as the endogenous control. Relative gene expression (fold change) were calculated using the 2−ΔΔCt method. To assess the effect of sublancin alone, expression in the sublancin-treated group (ii) was normalized to that of the untreated control (i). To evaluate the combinatorial effect, expression in the combination group (iv) was normalized to that of the vancomycin-alone group (iii) and that of the untreated control (i), respectively.

### Animals and ethics

Female C57BL/6 mice (6–8 weeks old, 20 g) were housed with sterilized food and water. All procedures were approved by the Institutional Animal Care and Use Committee.

### *In vivo* inntestinal VRE decolonization analysis

Intestinal colonization with VRE HN1 in mice was established according to previous published protocols (27), with minor modifications. Sepcially, mice were inoculated via oral gavage with VRE strain HN1 at a dose of 10^8^ CFU in 200 µL PBS.

After 7 days of colonization, each group animals were treated via oral gavage with vancomycin (20 mg/kg), vancomycin (20 mg/kg) + sublancin (20 mg/kg), vancomycin (20 mg/kg) + sublancin (20 mg/kg), sublancin (20 mg/kg) set PBS and linezolid as negative control and positive control, respectively. Each group were treated by oral gavage for 8 consecutive days. Fecal VRE loads were determined at baseline (day 0) and on days 3, 5, and 7 post-treatment initiation. Samples were collected, and diluted at rate 100mg/kg in PBS, then plated on BHA with vancomycin 80 mg/L, results are expressed as CFU/g feces (mean ± SD). On day 9, mice were euthanized, and VRE burdens in cecal and ileal tissues were quantified.

Statistical analysis of colony-forming unit (CFU) data was performed using two-way analysis of variance (ANOVA), followed by Dunnett’s post hoc test for multiple comparisons. In the figures, the asterisk (*) denotes significant differences between the indicated treatment groups and the linezolid-treated (positive control) group. Significance levels are denoted as follows: \**p* < 0.05, \*\**p* < 0.01, \*\*\*\**p* < 0.0001.

### Efficacy analysis in a *Galleria mellonella* infection model

The *in vivo* activity of sublancin in combination with vancomycin, and vancomycin alone, were evaluated in *Galleria mellonella larval* model. Larvae (250–350 mg; Huiyude Biotech) were randomly assigned into groups (n = 10 per group) and infected via injection of 10 μL containing 1.0 × 10⁷ CFU of a VRE HN1 strain. After 2 hours, each group were injected at the left posterior proleg treated, with PBS control ( vehicle control ), vancomycin ( 10 mg/kg ), sublancin ( 10 mg/kg ), and sublancin ( 10 mg/kg ) in combination with vancomycin groups ( 10+10 mg/kg ). Survival rate was monitored every 24 h up to 120 h.

For quantitative bacterial load analysis, larvae ( n=8 per group ) were randomly selected. Owing to the high mortality rate in the PBS group, eight dead larvae were randomly collected at 24 h intervals and homogenized. Serial dilutions were plated on enterococcus chromogenic medium (Beijing Land Bridge Technology) with 8 µg/mL vancomycin, and colonies were counted.

Statistical significance for survival data was determined by the log-rank (Mantel-Cox) test. CFU data were analyzed using one-way analysis of variance (ANOVA), with multiple comparisons against the control group performed by Dunnett’s post hoc test. Significance levels are denoted as follows: \**p* < 0.05, \*\**p* < 0.01, **** *p* < 0.0001 vs. control.

## RESULTS

### Antibacterial activity of sublancin and its potentiation of glycopeptide antibiotics againsr VRE *in vitro*

Sublancin exhibited intrinsic antibacterial activity against Gram-positive pathogens, including MRSA1, *poxtA and fexB* positive *E. faecalis* E033, E035 and VRE HN1, and its MICs were 12.5 mg/L, 20 mg/L, 80 mg/L, 160 mg/L.

Evaluation of checkerboard assays among combinations of sublancin with various beta-lactams, macrolides, or amphenicols showed no synergistic interaction (data not shown), while, sublancin strongly synergized with vancomycin against VRE HN1 (FIC index ≤ 0.5). This synergy indicates that sublancin specifically potentiates glycopeptide activity and is not a general adjuvant against other antibiotic classes.

To comfirm the efficacy, high-concentraction combination of sublancin and vancomycin resulted in clear culture broth with no bacterial growth of VRE after 24 h (Fig. 1A).

**Fig 1.**
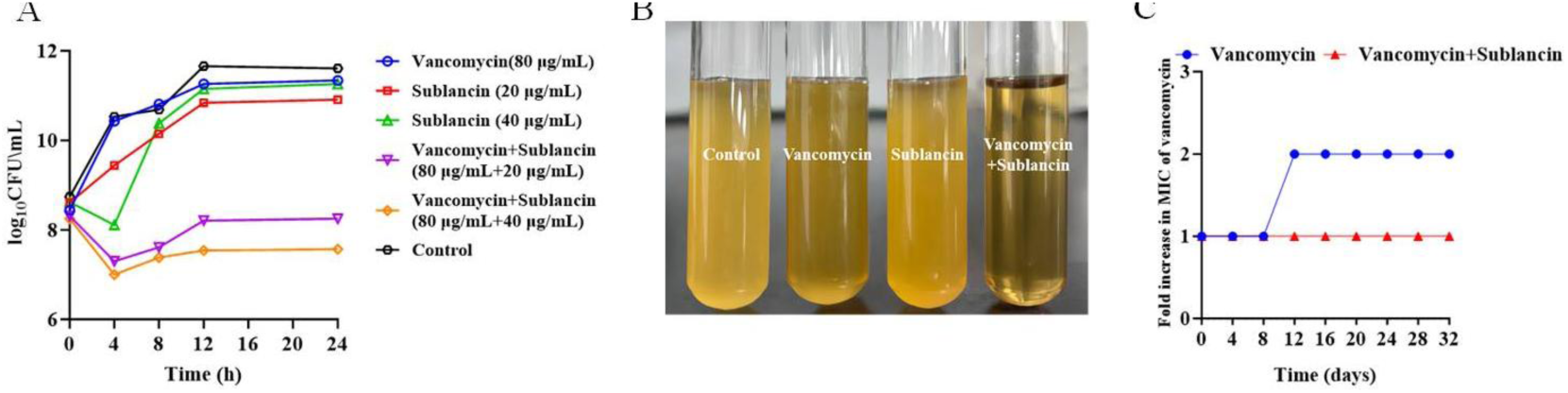
*In vitro* synergistic activity sublancin in combination with vancomycin against VRE HN1, as determined by time-killing assay (A).The visual and direct effect of sublancin in combination with vancomycin against the bacteria VRE HNl (B). Suppression of resistance development in VRE HN1 by sublancin in combination with vancomycin (C).

These findings indicate that sublancin possesses antibacterial activity against Gram-positive bacteria, and its most pronounced effect is the restoration of vancomycin susceptibility in VRE through synergistic interaction.

### Sublancin enhances vancomcyin efficacy and suppresses resistance development *in vitro*

Time-kill assay demonstrated that sublancin exhibits concentration-dependent bactericidal activity against VRE, achieving a ≥3-log10 CFU/mL reduction at 4× MIC and complete eradication within 24 h ( Fig. 1B ). In combination with vancomycin, sublancin significantly enhanced bacterial killing, producing a >3-log10 reduction in 24 h at subinhibitory concentrations.

A serial passage experiment revealed that the presence of sublancin prevented the emergence of higher MIC vancomycin during 32-day exposure to sub-MIC vancomycin and sublancin, whereas vancomycin alone produces high-resistant strains with 2-fold increase of MIC ( Fig. 1C ). Additionally, sublancin long exposure to did not induce resistance in VRE HN1, *E. faecalis* E035 over 90 days.

Collectively, these findings indicate that sublancin not only potentiates the bactericidal activity of vancomycin, but also suppresses the development of glycopeptide resistance *in vitro*.

### Sublancin enhances vancomcyin efficacy *in vivo*

Following the demonstration of *in vitro* activity against VRE HN1, the efficacy of sublancin was further evaluated using a *Galleria mellonella* infection models and murine decolonization assays.

In the *G. mellonella* model, treatment with the combination of vancomycin (10 mg/kg) and sublancin (10 mg/kg) significantly increased larval survival to 80% at 96 h post-infection, compared to 20% in the PBS control group ( *p* < 0.001, Fig. 2A ). the combination of vancomycin (10 mg/kg) and sublancin (10 mg/kg) also reduced bacterial loads in larval, achieving a 6.19-log10 CFU/mL decrease in VRE HN1 counts at 24 h relative to the control ( *p* < 0.001, Fig. 2A ). Sublancin exhibited dose-dependent protection against HN1 in *G. mellonella*. Larvae administered sublancin (10 mg/kg) showed an 50% survival rate at 72 h post-infection, compared to 20% in PBS-treated controls ( *p* < 0.001, Fig. 2B ). This survival benefit correlated with a 6.19-log10 reduction in bacterial burden (Fig. 2B). No toxicity was observed at therapeutic doses, as evidenced by 96% survival in uninfected larvae.

**Fig 2.**
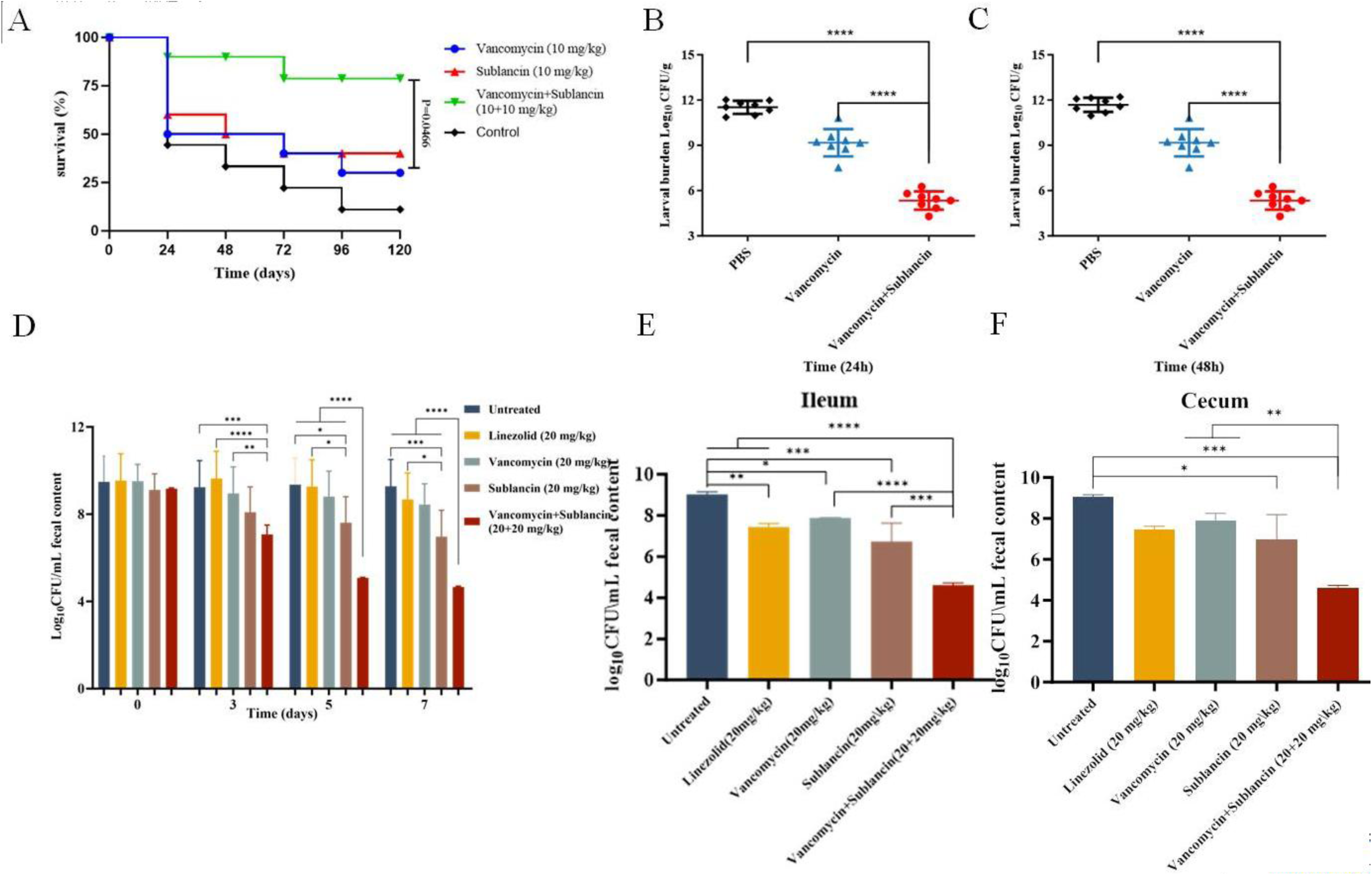
Efficacy of sublancin in combination with vancomycin in a *Galleria mellonella* infection model and a mouse decolonization model. (A-C) *Galleria mellonella* infection model. Larvae were infected with VRE strain HN1, and after 1 h, administered a single injection of vancomycin alone, sublancin alone, or a combination of both in. (A) Survival rates were monitored over 5 days (Kaplan-Meier analysis). Bacterial loads in larvae were quantified at (B) 24 h and (C) 48 h post-treatment. (D-F) Mouse decolonization model. Mice were colonized with VRE HN1. Fecal bacterial loads were monitored on days 0, 3, 5, and 7 (D) to assess decolonization dynamics. Untreated and linezolid-treated groups served as the negative and positive controls, respectively. Following 8 consecutive days of dosing with each antibiotic or their combination), bacterial loads in the (E) ileum and (F) cecum were determined. Data are expressed as mean CFU ±SD. Statistical analysis was performed using one-way ANOVA followed by Dunnett’s multiple-comparison test (\**p* < 0.05, \*\**p* < 0.01, ** ** *p* < 0.0001 vs. control).

In a murine gut decolonization model of VRE strain HN1, sublancin monotherapy and the sublancin-vancomycin combination significantly reduced the fecal VRE HN1 load compared to both vehicle and linezolid control groups ( Fig. 2C ). By day 3, these treatments produced at least a 1-log₁₀ CFU reduction relative to the vehicle, while linezolid showed minimal effect. Continued treatment with sublancin or the combination yielded sustained decolonization, with reductions ranging from 1.74 to 4.47 log₁₀ 98.2–99.9% reduction by day 5 ( *p* < 0.0001 ), significantly greater than the 0.55-log₁₀ CFU/g 71.0% reduction observed with vancomycin alone ( *p* < 0.0001, Fig. 2D ). In contrast, linezolid achieved only a 0.09-log₁₀ 19% reduction ( *p* < 0.0001 ).

The vancomycin - sublancin combination further reduced the VRE HN1 burden by 2.31–4.62-log₁₀ ( corresponding to 99.5–99.9% reduction ) by day 7 (*p* < 0.0001 ). Bacterial loads in ileal and cecal tissues were also markedly decreased: sublancin alone reduced counts by 2.06–2.28-log₁₀ ( 98.7–99.5% reduction ), whereas the combination achieved reductions of 4.42–4.56-log₁₀ ( >99.9% reduction, *p* < 0.01, Fig. 2E ). Collectively, the vancomycin - sublancin regimen demonstrated the highest efficacy, approaching complete bacterial eradication.

### Sublancin affectes bacterial physiological state and energy metabolism

The effect of sublancin on membrane permeability was assessed using fluorescent probes. The permeability of OM was monitored using NPN probe, which fluoresces upon intercalation into the phospholipid bilayer. In VRE strain HN1, it was observed that sublancin induced a dose-dependent increase in NPN fluorescence, indicating that sublancin increased OM permeability ( Fig. 3A ). This disruption of the OM is consistent with its role in potentiating vancomycin activity against this strain. In contrast, PI assay revealed no significant increase in fluorescence, suggesting that sublancin exposure did not immediately influenced cytoplasmic IM integrity. Membrane depolarization was evaluated using the DiSC_3_(5). Treatment with sublancin resulted in a significant decrease in fluorescence, consistent with membrane depolarization ( Fig. 3B ). To assess the influence to energy metabolism, extracellular ATP, intracellular ATP and total ATP levels were quantified. Intracellular and total ATP were significantly depleted in sublancin-treated group, while extracellular ATP increased correspondly ( Fig. 3C ). these results may indicate impaired energy metabolism—possible through disruption of substrate level phosphorylation, oxidative phosphorylation, or both- and suggest possible membrane leakage. The result of depolarization and ATP depletion are likely to further compromise oxidative phosphorylation.

**Fig 3.**
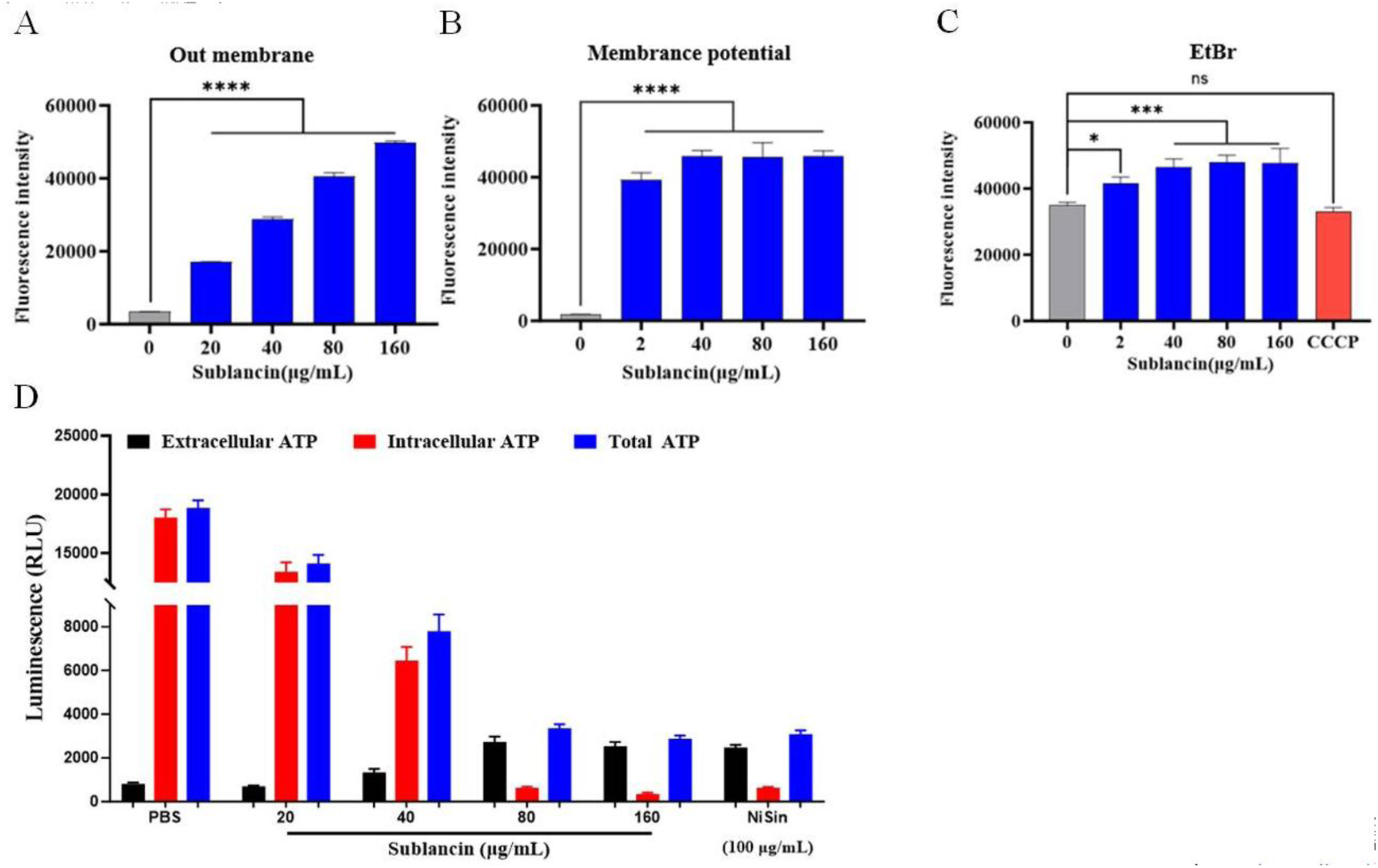
Potential mechanism of sublancin in combination with vancomycin. (A) Effect of increasing sublancin concentrations on the integerity of bacterial outer membrane in VRE HN1. (B) Alteration of membrane potential in VRE HN1 following sublancin treatment.(C) Levels of extracellular, intracellular, and total ATP in VRE HN1 after exposure to sublancin. Exposure to increasing concentraction of sublancin led to decreased intracellular and total ATP, alongside a slight increase in extracellular ATP. Data are mean ± SD. Significance was assessed by nonparametric one-way ANOVA (ns, not significant, \**p* < 0.05, \*\*\**p* < 0.001, \*\*\*\**p* < 0.0001).

Notably, no concurrent increase in the PMF or reactive oxygen species (ROS) was detected. Efflux pump activity, assessed using EtBr accumulation assay, was not influenced by sublancin exposure ( data not shown ).

### Sublancin modulates transcription of the *vanA* operon

To elucidate the mechanism underlying the synergy activity, we quantified the impact of sublancin on *vanA* operon transcriptionvia qRT-PCR gene expression was analyzed under three conditions: (1) the effect of sublancin alone, where the sublancin-treated group was normalized to the untreated control, (2) the combinatory effect, where the sublancin-plus-vancomycin group was normalized to the vancomycin-alone group, and (3) the combinatory effect, where the sublancin-plus-vancomycin group was normalized to untreated control. The results are presented in Fig. 4.

**Fig 4.**
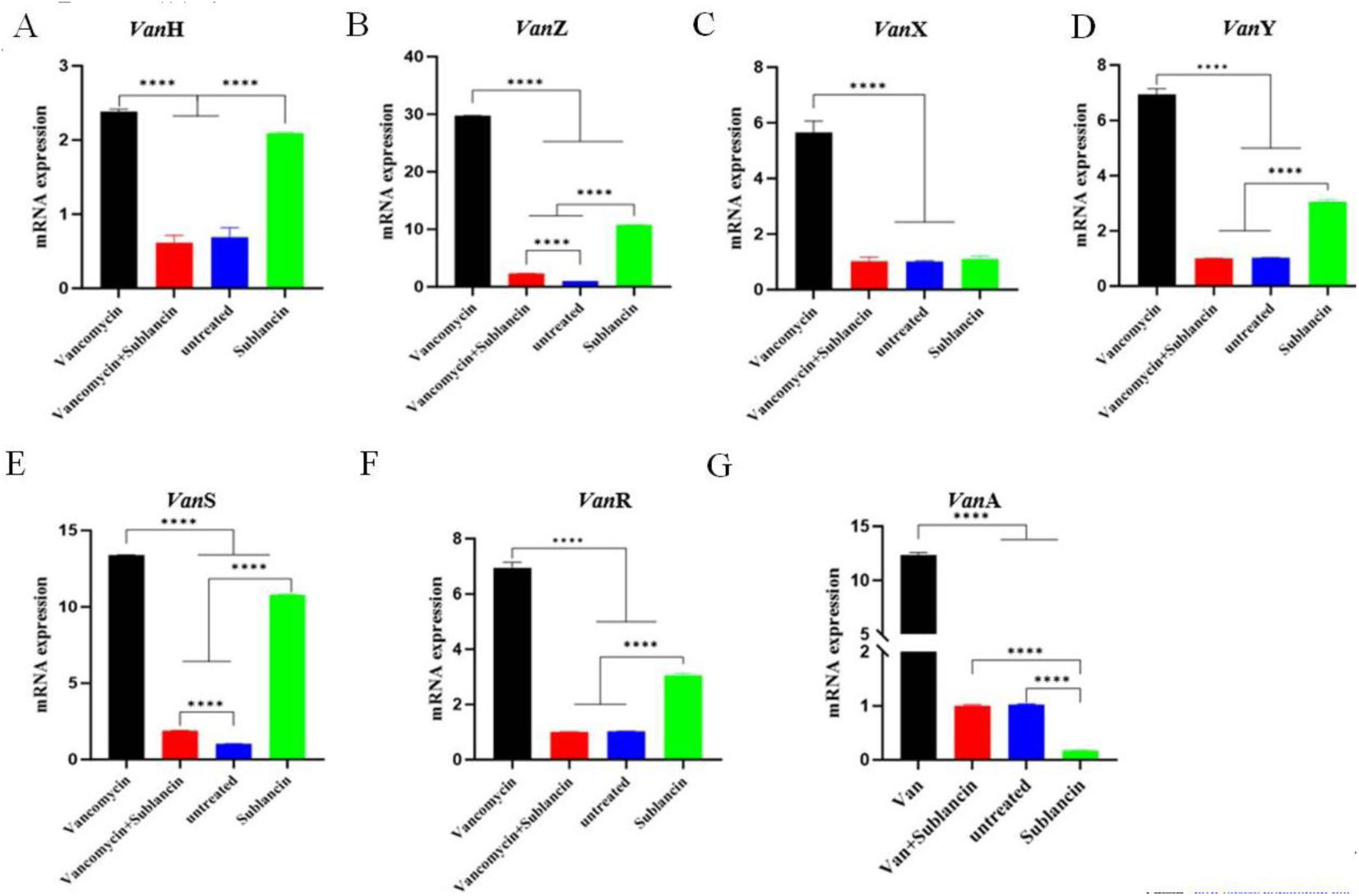
Transcription modulation of the *vanA* operon by sublancin alone and sublancin in combination with vancomycin. The *vanA* operon was assessed under four conditions: untreated (negative conrrol), vancomycin alone(van), sublancin alone, and the combination of vancomycin and sublancin. Relative transcript levels of the following genes are shown: *vanH* (A ), *vanZ*(B), *vanX* (C), *vanY*(D), *vanS* (E),*vanR*(F) and *vanA* (G). Data are presented as mean ±standard deviation (SD). Significance levels are denoted as follows: \*\*\*\**p* < 0.0001.

Exposure to a subinhibitory concentration of sublancin (20 µg/mL) for 12 h, altered the transcription profile of the *vanA* operon. The regulatory genes *vanR*, *vanS*, *vanH*, *vanX* and *vanZ* were significantly up-regulated by 298.3%, 1047.6%, 303.3%, 108.6%and 1066.3%, respectively, compared with the untreated control. In contrast, the expression of the structrural/ resistance gene *vanA* and *vanY* was down-regulated to 16.7% and 78.8% of the untreated control levels, with *vanA* the most significant suppression.

Further, when combined with vancomycin, sublancin dramatically attenuated the antibiotic - induced expression of all tested *vanA* operon genes. Relative to the vancomycin-alone group, the transcript levels of *vanR*, *vanS, vanH*, *vanA*, *vanX, vanY,* and *vanZ* were reduced to 14.1%, 14.4%, 25.7%, 8.1%, 18.1%, 18.0% and 7.8%, respectively. Compared to the baseline (untreated control), the expression of *vanR*, *vanH*, and *vanA* were reduced to 97.7% , 88.8%, and 97.7% of control levels, respectively.

## DISCUSSION

Sublancin was initially identified from *Bacillus subtilis* 168 in the late 1970s as an antimicrobial substance, exhibiting activity against other Gram-positive bacteria, including Bacillus and Staphylococcus species, along with significant immunomodulatory properties (23, 28). For decades, its post-translational modifications and activity spectrum led to its classification as a member of the lantibiotic family—ribosomally synthesized and post-translationally modified peptides (RiPPs) characterized by thioether cross-links formed by lanthionine and methyllanthionine residues (29). However, a pivotal structural study in 2011 redefined sublancin not as a lantibiotic, but as the prototype of a novel class of S-linked glycopeptides (30).

Beyond its direct antimicrobial effects, sublancin exhibits a broad spectrum of biological functions, including bactericidal activity and sophisticated regulatory roles within bacterial communities. Prior to this study, sublancin was known to exert potent, direct bactericidal activity primarily against other Gram-positive bacteria, with its mechanism dependent on the bacterial phosphoenolpyruvate:sugar phosphotransferase system (PTS) (31, 32). In the present work, we demonstrate that sublancin potentiates vancomycin activity against *vanA*-type VRE through a specific mechanism involving transcriptional repression of resistance determinants.

The key finding of this study is that sublancin at a subinhibitory concentration, differentially modulates transcription of the *vanA* operon, both alone and combination with vancomycin. Unlike SLAY-P1 (24), which did not significantly alter operon expression, sublancin alone markedly upregulated the sensor-regulator genes *vanS* and *vanR* while suppressing the core resistance genes *vanH*, *vanA*, and *vanX*. More importantly, the combination of sublancin with vancomycin resulted in a profound, synergistic repression of the entire operon—an effect similarly observed with the SLAY-P1–vancomycin combination.

Concurrently, sublancin compromises the physical barrier that limits vancomycin access. We found that sublancin increases outer membrane permeability in a dose-dependent manner, which is crucial for gram-positive bacteria like enterococci that can exhibit reduced drug uptake. Furthermore, sublancin induces cytoplasmic membrane depolarization and a severe depletion of intracellular ATP, indicating a collapse of central bioenergetic processes. The resulting loss of membrane potential and energy charge would be expected to inhibit the ATP-dependent ligase activity of *VanA*, thereby synergizing with the transcriptional suppression to comprehensively disable the resistance pathway.

Thus, we propose a model in which sublancin acts on two fronts: it disarms the bacterium by suppressing the expression of the inducible resistance machinery, while simultaneously weakening cell defenses through disruption membrane integrity and energy metabolism. This multifaceted action renders VRE susceptible to vancomycin, overcoming a key mechanism of high-level resistance. These findings position sublancin, and potentially similar bacteriocins, as promising adjuvants for revitalizing glycopeptide antibiotics against resistant pathogens.

## ACKNOWLEDGMENTS

We received no financial support for the research, authorship, or publication of this article.

## Supplemental Material

**Table S1.**
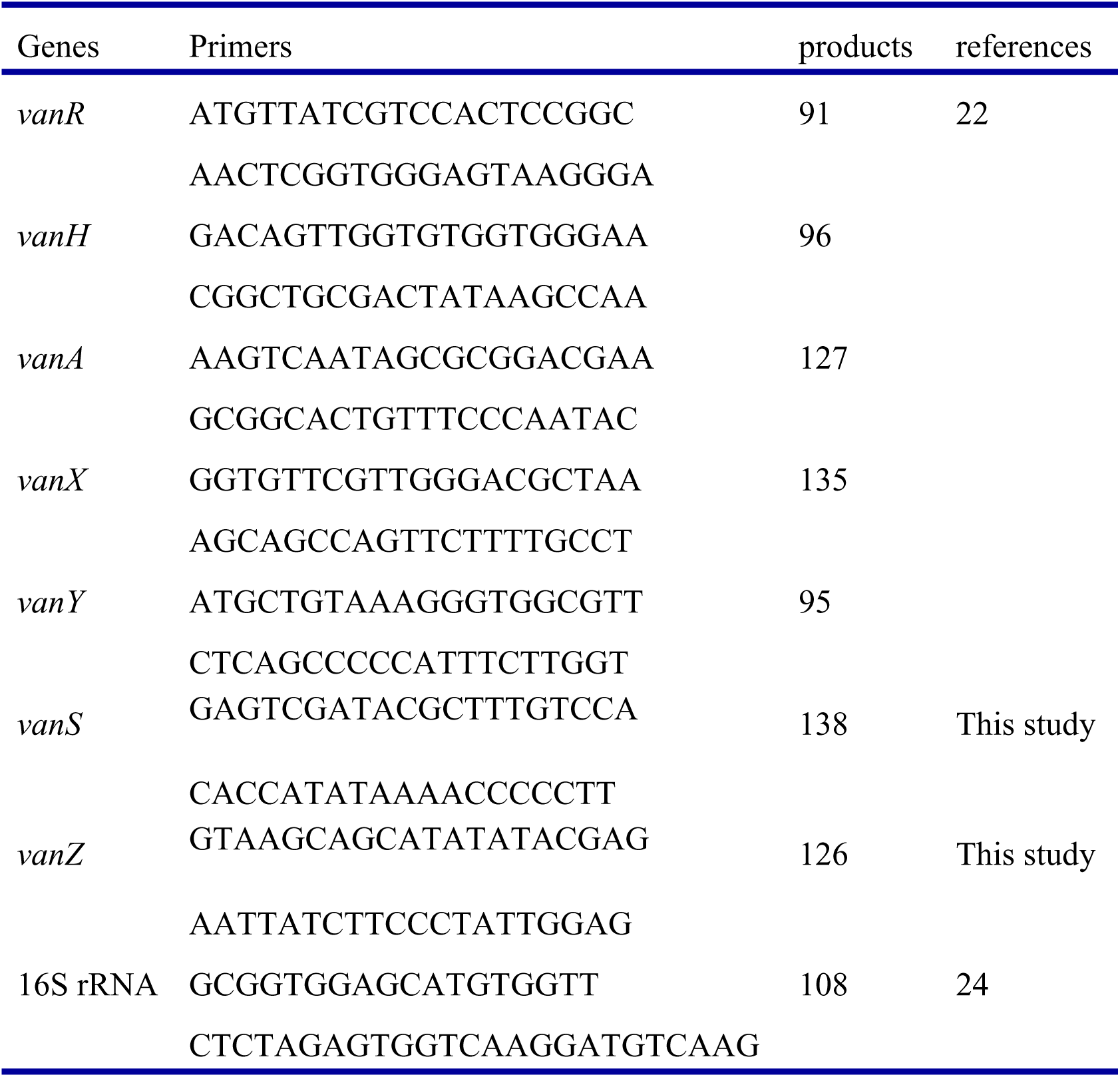
Basic information of RT-PCR Primers for the *VanA* Operon.

